# Phenotypic analysis of Arabidopsis transgenic plants constitutively expressing the P6 protein from *Cauliflower mosaic virus* or mutant alleles thereof

**DOI:** 10.1101/023283

**Authors:** Angèle Geldreich, Christophe Himber, Gabrielle Haas, Mario Keller, Olivier Voinnet

## Abstract

This laboratory has previously shown that nuclear import of CaMV P6 is required for infection and suppression of the RNA silencing factor DRB4. EMBO J., 27:2102 - 2112. In the original paper, there were inadvertent errors in the mounting of a figure. We have now repeated the experiments concerned, using the seed stock of the corresponding transgenic plants. The results presented here confirm those reported in the original paper. The plants expressing P6 are strongly stunted with chlorotic and serrated leaves. Conversely, the transgenic expression of various P6 mutant alleles do not induce this phenotype.

## Introduction

The P6 protein encoded by *Cauliflower mosaic virus* (CaMV) is a symptom and host range determinant known to suppress transgene silencing (Love et al, 2007). P6 is also essential for CaMV replication and particle assembly in single cells, where it aggregates into amorphous cytoplasmic bodies termed viroplasms (Kobayashi *et al*., 1998; Haas *et al*., 2005). This prevalent cytoplasmic distribution is contributed by the N-terminal domain A (spanning aa 1-111), which contains a nuclear export signal (NES) overlapping the residues required for P6–P6 intermolecular interactions and viroplasm formation. Nonetheless, a small fraction of P6 is also found in the nucleus but its mode of nuclear entry and potential functions in this organelle had remained largely unknown (Haas *et al*., 2005).

## Summary

We showed in Haas *et al.* (2008) that monomeric P6 is imported into the nucleus through two importin-α-dependent nuclear localization signals: a conventional bipartite basic motif spanning amino acids (aa) 314-328 referred to as ‘NLS’, and a non-conventional motif comprising aa 219-234 referred to as NLSa. We also showed that nuclear import enables P6 to suppress RNA silencing in several reporter systems in Arabidopsis. Silencing suppression relies, at least in part, upon the interaction of P6 with the double-stranded (ds)RNA binding protein DRB4, which is required for optimal activity of the major Arabidopsis antiviral protein, Dicer-like 4 (DCL4); DCL4 produces small interfering (si)RNAs from viral RNA with double-stranded features (reviewed in Ding and Voinnet, 2007). DRB4 is also required for optimal accumulation of so called trans-acting (tasi) RNAs, which are endogenous products of DCL4 involved in leaf patterning and juvenile-to-adult phase transition in Arabidopsis (Yoshikawa *et al*., 2005; Adenot *et al*., 2006). A dsRNA-binding domain contained within the so-called “mini-TAV” region of P6 (spanning aa 111-242) was found dispensable for DRB4 inhibition and RNA silencing suppression by P6 (Haas *et al.,* 2008).

The above findings relied, in part, on the use of transgenic Arabidopsis expressing the wild type P6 protein or several of its derivatives carrying point mutations or deletions in its key domains. These included mutant P6Δa, in which the non-conventional NLSa is deleted, P6ΔNLSΔa, in which the two NLS are both deleted, P6_m3_ in which the EKI tripeptide contained in the NES motif is mutated, and P6ΔdsR in which the dsRNA-binding motif of the mini TAV region is deleted. Several independent transgenic lines were generated for each P6 mutant, and plants expressing similar levels of protein were selected for molecular analyses, including those of endogenous tasiRNAs depicted in (Haas *et al*. 2008; Figure 4E).

Figure 4A-D of Haas et al. (2008) was intended to compare the phenotypes of the P6 transgenic plants and of the plants expressing the above-mentioned P6 variants. As previously reported, the P6-expressing plants were very stunted and displayed strongly serrated and chlorotic leaves; in contrast, none of the plants expressing the P6 mutant variants had an overt phenotype compared to non-transgenic, wild type Arabidopsis. However, it was brought to our attention that two of the four images depicted in the original panel 4D (P6_m3_ and P6ΔdsRNA) were erroneous duplications of images found in other, unrelated figure panels of Haas *et al*. (2008).

Although we have confirmed that these duplications occurred inadvertently at the figure mounting stage, we were unfortunately unable to retrieve the source data for the images depicted in panel 4D. In order to dispel any doubts on the original results, the seed stock of the corresponding transgenic plants was retrieved and plants grown side-by-side under identical, controlled conditions (see material and methods). Pictures of whole plants were taken 5 weeks post-germination (wpg) and are displayed in Figure 1. Specific sections and leaves of the various plants are shown in Figure 2. These images confirm that P6-expressing plants are strongly stunted with serrated and chlorotic leaves (Figure 1 and Figure 2B-C) and that, by contrast, the plants and leaves expressing the various P6 mutant alleles are not distinctively different from non-transgenic, wild-type Arabidopsis (Figure 1 and Figure 2D).

**Figure 1.**
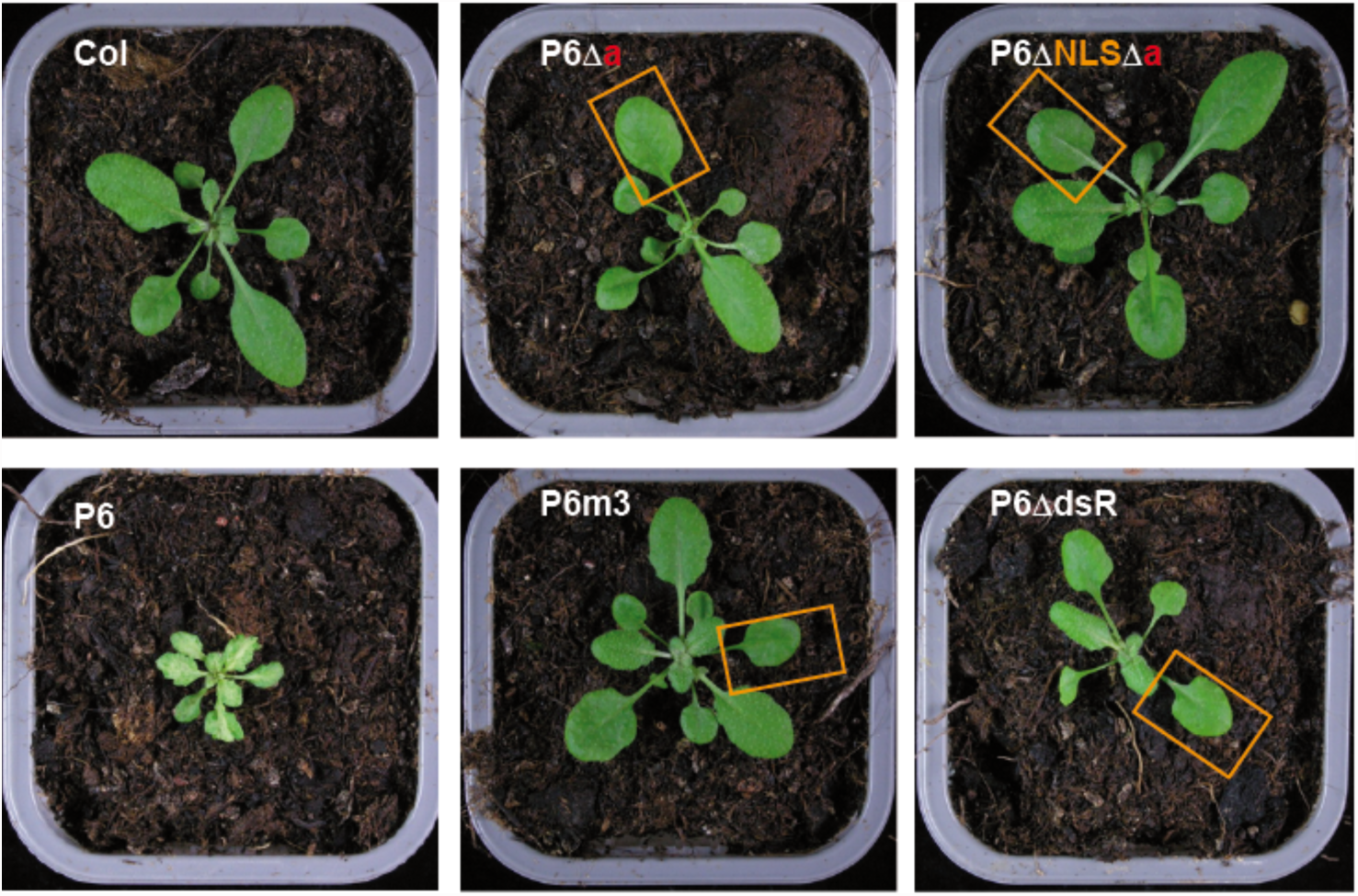
P6-expressing transgenic plants in Arabidopsis ecotype Col-0. Transgenic expression of the CaMV P6 protein induces stunting of the plant, chlorosis and serration of the leaf (P6) compared to the wild-type Arabidopsis plant (col). By contrast, none of the plants expressing the P6 deletion mutants (P6Δa, P6ΔaΔNLS or P6ΔdsR) or punctual mutant (P6m3) exhibit an overt phenotype compared to the control plant (Col). The orange squares highlight the leaves used for the magnified views presented in Figure 2.

**Figure 2.**
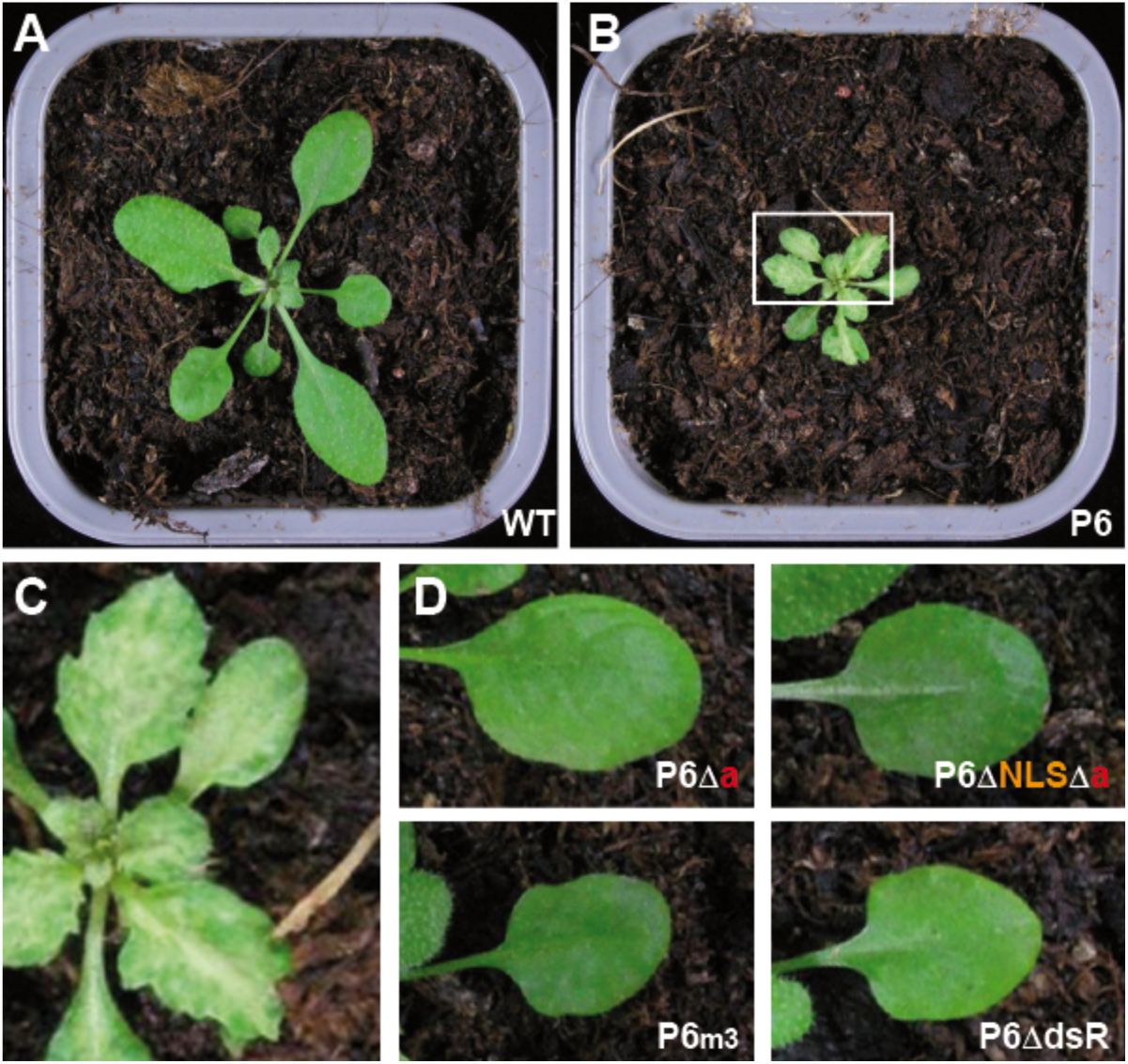
Phenotypic comparison of the leaves produces by P6 transgenic plants. The wild-type, non-transgenic (WT) Arabidopsis ecotype Col-0 (A) and the P6-transgenic plant (B) are the same as presented in Figure 1. The white square on the P6 transgenic plant in (B) serves to highlight the area selected for the magnified view of P6-induced symptoms (chlorosis, leaf serration) presented in (C.) (D) None of the P6 mutants induces such leaf anomalies when transgenically expressed in the Col-0 ecotype. The leaves were magnified from the pictures presented in Figure 1 and were selected according to the phyllotaxy.

## Material and methods

The P6 transgenic Arabidopsis and Arabidopsis expressing the P6 mutant variants were described in Haas *et al.* (2008). Sterilized seeds were initially sown *in vitro* on selective medium (MS salts Sigma M5519 + 0,5g/L MES Duchefa + 1% Saccharose Duchefa + 0,8 % Agar Agar Duchefa, pH 5,7), supplemented with Kanamycine (Duchefa) at a final concentration of 50 mg/L, except for control wild type plants. *In vitro* growth conditions were: Day/Night 16/8, Temperature 21/17, Lights: 4x T8 BioLux 58W Intensity. Seventeen days after germination, seedlings were pricked out and transplanted into soil in individual pots (LAT Terra Standard supplemented with Basacote Tabs (11-11-18) 2g/L). Soil growth conditions were set in dedicated growth cabinets: Day/Night 12/12, Temperature 21/18, Lights type 4x T8 BioLux 58W Intensity. Representative whole plants were imaged 34 days post-germination, using a handheld Nikon CoolPix camera.

